# A nuclear pore sub-complex restricts the propagation of Ty retrotransposons by limiting their transcription

**DOI:** 10.1101/2021.01.05.425438

**Authors:** Amandine Bonnet, Carole Chaput, Benoit Palancade, Pascale Lesage

**Affiliations:** Université de Paris, Institut de Recherche Saint-Louis, INSERM U944, CNRS UMR 7212, F-75010 Paris, France; Université de Paris, CNRS, Institut Jacques Monod, F-75006 Paris, France

**Keywords:** Ty LTR-retrotransposons, nuclear pore complexes, Nup84 complex, yeast

## Abstract

Beyond their canonical function in nucleocytoplasmic exchanges, nuclear pore complexes (NPCs) regulate the expression of protein-coding genes. Here, we have implemented transcriptomic and molecular methods to specifically address the impact of the NPC on retroelements, which are present in multiple copies in genomes. We report a novel function for the Nup84 complex, a core NPC building block, in specifically restricting the transcription of LTR-retrotransposons in yeast. Nup84 complex-dependent repression impacts both *Copia* and *Gypsy* Ty LTR-retrotransposons, all over the *S. cerevisiae* genome. Mechanistically, the Nup84 complex restricts the transcription of Ty1, the most active yeast retrotransposon, through the tethering of the SUMO-deconjugating enzyme Ulp1 to NPCs. Strikingly, the modest accumulation of Ty1 RNAs caused by Nup84 complex loss-of-function is sufficient to trigger an important increase of Ty1 cDNA levels, resulting in massive Ty1 retrotransposition. Altogether, our studies expand our understanding of the complex interactions between retrotransposons and the NPC, and highlight the importance for the cells to keep retrotransposon under tight transcriptional control.

**AUTHOR SUMMARY:** Retroelements, which replicate by reverse transcription of their RNA into a cDNA that is integrated into the host genetic material, play an important role in the plasticity of eukaryotic genomes. The study of yeast retrotransposons has led to the identification of host factors that limit retroelement mobility, including components of the nuclear pore complex (NPC), most of them still awaiting mechanistic characterization. Here, we investigated the contribution of the Nup84 complex, a core NPC scaffold, to retrotransposon biology in budding yeast. Our findings uncover that the Nup84 complex restricts the transcription of phylogenetically-distinct Ty retroelements. By focusing on Ty1 retrotransposons, we provide evidence that repression by the Nup84 complex depends on the maintenance at the NPC of the SUMO-protease Ulp1, an essential enzyme of the SUMO pathway with multiple targets in the transcription machinery. We finally show that this transcriptional control is critical for genome dynamics, since a small increase in the accumulation of Ty1 RNAs leads to massive retrotransposition. Our data reveal that although relatively abundant, Ty transcripts are limiting for retrotransposition, underscoring the importance of a tight control of their expression. They also characterize a new non-canonical function of NPCs, confirming their connection with genome expression and stability.

## INTRODUCTION

As the unique gates between the cytoplasm and the nucleus, nuclear pore complexes (NPCs) are key components of eukaryotic cells. NPCs are modular, megadalton-sized multiprotein assemblies built from multiple copies of ~30 distinct proteins called nucleoporins (Nups) and associated into sub-complexes (reviewed in [1,2]). The overall structural organization of the NPC is conserved across eukaryotes, with a core scaffold embedded within the nuclear envelope and peripheral components, namely the cytoplasmic filaments and the nuclear basket, extending towards the cytoplasm and the nucleoplasm, respectively.

Beyond their canonical role in the selective transport of proteins and RNAs, NPCs have also emerged as key platforms for the three-dimensional organization of the genome, thereby impacting the regulation of gene expression and the maintenance of genetic integrity (reviewed in [3–5]). Genome-wide approaches, together with the live imaging of individually-tagged loci, have revealed that nucleoporins associate mostly with transcriptionally-active genes, but also with repressed regions and chromatin boundaries, from yeasts to metazoans [6–14]. While a subset of nucleoporins can bind their target sequences away from the NPC [15–17], these interactions typically occur at the nuclear envelope, as exemplified by the well-described recruitment of induced genes to the pores in budding yeast [18–21]. Such functional associations between NPCs and the genome are relevant for the establishment of expression patterns, as revealed by the transcriptional changes triggered in *cis* of the Nup-interaction sites upon nucleoporin loss-of-function (reviewed in [5,22]). Although NPCs have been proposed to serve as scaffolds for transcriptional activities, impacting chromatin organization and mRNA biogenesis [5,20], the direct and indirect mechanisms underlying their function in gene expression remain to be fully elucidated.

While a critical role in transcription has been assigned to components of the nuclear pore basket, which are closest to the the genome [11,23–26], gene expression has also been reported to depend on the core NPC scaffold [9,27,28], including the Y-complex (a.k.a. the Nup84 complex in budding yeast). Apart from its critical role in NPC distribution and mRNA export (reviewed in [29]), the yeast Y-complex has been shown to participate to the maintenance of genome integrity [27,30,31] and to multiple steps in transcriptional regulation, including activation, repression and elongation [31–35]. In some cases, the Y-complex was shown to regulate these processes by tethering the SUMO-deconjugating enzyme Ulp1, an essential member of the SUMO modification pathway with multiple targets in the DNA and RNA metabolism machineries (reviewed in [36–38]).

So far, most studies exploring the role of the NPC in genomic regulations focused on protein-coding genes, with few reports indicating its function at other loci (e.g. RNA polymerase III-transcribed genes [39,40]). In line with the difficulties associated with the analysis of repetitive sequences [41], little is known about the influence of NPCs on the RNA polymerase II (RNAP II)-dependent transcription of transposable elements (TEs). TEs are ubiquitous in eukaryotes and represent a significant fraction of their genomes (reviewed in [42]). The *S. cerevisiae* genome hosts five families of LTR-retrotransposons (Ty1 to Ty5) harboring the same basic genomic organization, consisting of two direct terminal repeats (LTRs) flanking the *GAG/TYA* and *POL/TYB* open reading frames. Like their retrovirus counterparts, Ty retrotransposons replicate by reverse transcription of their RNA genome into a cDNA copy that is stably integrated into the host-cell genome by their self-encoded integrase. Except for Ty3, which is a *Metaviridae* (gypsy-like element), the other yeast Ty elements belong to the *Pseudoviridae* group (copia/Ty1 elements). Ty1 RNAs account for 0.1-0.8% of total cellular RNA [43,44] and thus represent an important part of the yeast transcriptome that remains moslty unexplored. Ty1 transcription depends on several transcription factors that bind to the Ty1 promoter (i.e., Gcr1, Ste12, Tec1, Mcm1, Tea1/Ibf1, Rap1, Gcn4, Mot3 and Tye7) or belong to chromatin-remodeling complexes (Swi/Snf, SAGA and ISWI) (reviewed in [45]). In addition, dozens of cellular factors identified as regulators of Ty1 replication (reviewed in [45]), including subunits of the Nup84 complex [46–49], still await mechanistic characterization.

In this study, we investigated the contribution of the Nup84 complex to the yeast *S. cerevisiae* transcriptome with a special focus on retrotransposon transcripts. Using a combination of genome-wide approaches and dedicated reporter systems, we show that the Nup84 complex represses the transcription of most copies of the Ty1, Ty2 and Ty3 retrotransposons. By genetically disentangling the multiple roles of the Nup84 complex, we further establish that its function in the tethering of the SUMO-protease Ulp1 to the NPC is essential for the transcriptional control of Ty1 retrotransposons. Finally, using a suite of tools to assess each stage of the Ty1 retrotransposition cycle, we demonstrate that, although the increase in Ty1 RNA levels appears minor upon loss-of-function of the Nup84 complex, it has major consequences on Ty1 retrotransposition. The Nup84 complex thus appears essential to keep retrotransposon mobility under control, highlighting the previously underestimated importance of fine-tuning retroelement transcription to limit their propagation in the yeast genome.

## RESULTS

### The Nup84 complex specifically restricts RNA accumulation of LTR-retrotransposons

To investigate the role of the Nup84 complex on gene expression, we performed a transcriptomic analysis in two different mutants deleted for non-essential subunits of this NPC subcomplex (*nup133*Δ and *nup84*Δ). To extend our analysis beyond protein-coding genes and include retrotransposon transcripts, which are mostly non-polyadenylated [50], RNA-seq was performed after depletion of the highly-abundant ribosomal RNAs but without selection for polyadenylated RNAs. Since Ty1 retrotransposition is optimal at 20-22°C [51] and most Nup84 complex mutants are thermosensitive (Fig S1A), all the experiments were further conducted at 25°C to guarantee both Ty1 mobility and the growth of mutant strains. Three independent cell cultures were analyzed for each genotype with a high level of reproducibility between samples of the same strain (Fig S1B). Globally, the mean transcriptomes of *nup133*Δ and *nup84*Δ cells barely differed from that of WT cells (Fig 1A). In these strains, 32 full-length copies of Ty1, 13 full-length copies of Ty2 and 2 full-length copies of Ty3 produce RNAs suitable for reverse transcription while the 3 copies of Ty4 only yields truncated transcripts lacking essential sequences for reverse transcription, and the only Ty5 copy is truncated and not expressed [52,53]. Since the sequences of the Ty copies within each family are very close, Ty RNAs are usually discarded from genomic analyses that focus on single-mapping reads. To circumvent this issue, we used multi-mapping reads, randomly assigned to these repeated sequences of the genome, in order to evaluate the bulk of Ty RNAs in WT and *nup* transcriptomes. In this way, we confirmed that Ty1 transcripts are among the most abundant RNAs in WT cells while Ty2 and Ty3 transcripts are less abundant, correlating with lower genomic copy number [53] (Fig 1B). Most importantly, our analysis revealed that Ty1, Ty2 and Ty3 transcripts are over-represented about 1.5-fold and 2-fold in the transcriptome of *nup133*Δ and *nup84*Δ mutants, respectively (Fig 1B). Differential expression analysis further confirmed an important increase in Ty1 and Ty3 RNA levels in both Nup84 complex mutants (Fig 1C), while Ty2 transcripts were mostly up-regulated in *nup84*Δ cells. Notably, these tendencies were not observed for mRNAs, tRNAs or other ncRNAs (Fig 1C), demonstrating that Nup133 and Nup84 specifically target LTR-retrotransposon transcripts.

**Figure 1:**
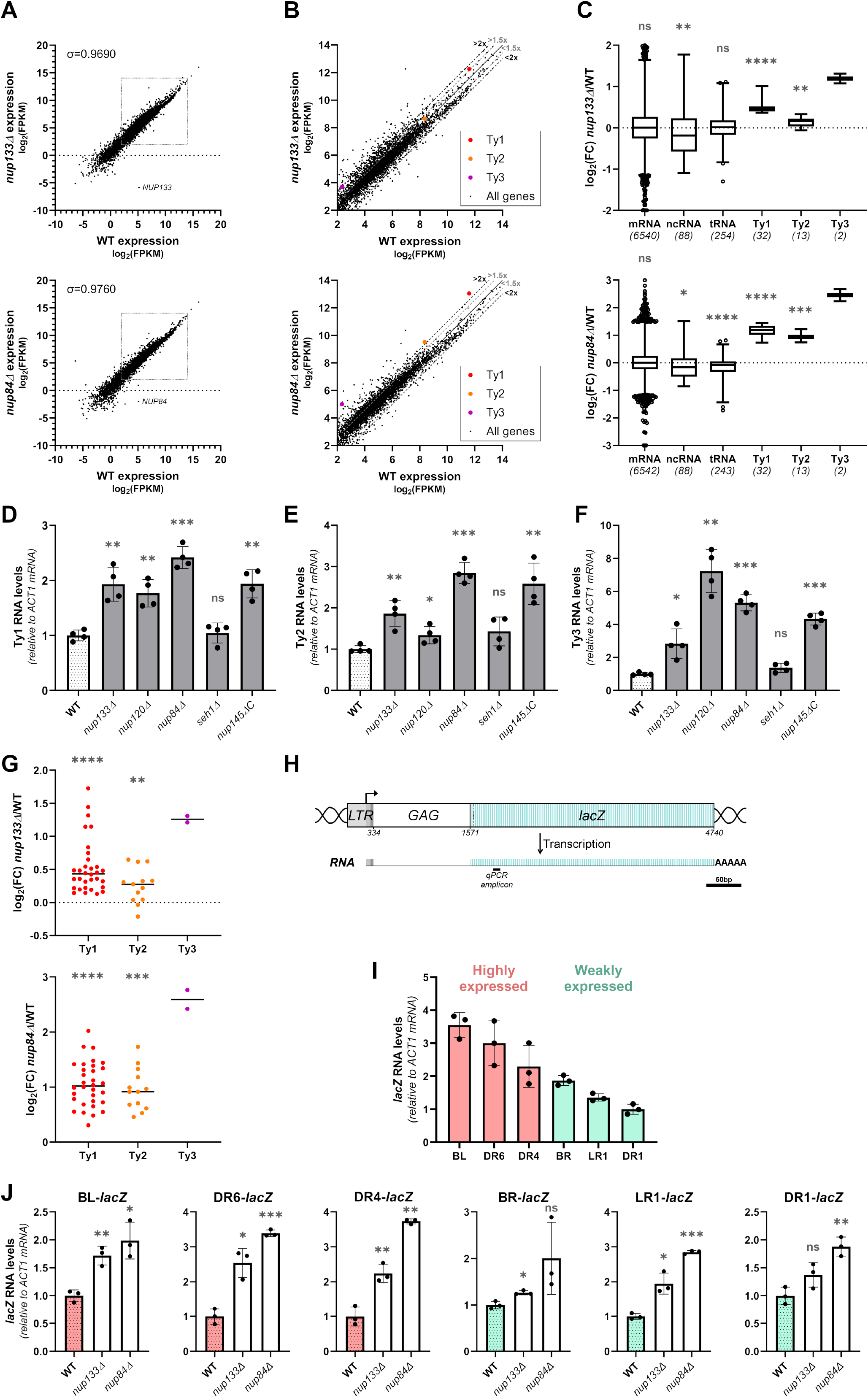
The Nup84 complex specifically restricts Ty RNA accumulation. **(A)** Scatterplots showing RNA abundance in *nup133*Δ (top) or *nup84*Δ (bottom) mutants vs WT cells based on RNA sequencing data. RNA levels are represented as log2 of fragments per kb per million reads mapped (FPKM). Each value is the average of three biological replicates per strain. The position of *NUP133* and *NUP84* transcripts are highlighted and the Spearman’s correlation coefficient is indicated. The framed area is zoomed in (**B**). **(B)** Scatterplots as described in (**A**) but restricted to the range of expression of Ty transcripts. Ty1, Ty2 and Ty3 RNA abundance is highlighted with red, orange and purple dots, respectively. The thresholds of 1.5 or 2-fold change between the 2 strains are indicated by grey or dark dashed lines, respectively. **(C)** Boxplot analysis of log2 fold change (FC) of different categories of transcripts in *nup133Δ (top panel)* or *nup84Δ (bottom panel)* mutants relative to WT cells. The number of transcripts expressed in both strains and considered in each category is indicated in brackets. **(D-F)** Ty1 **(D)**, Ty2 **(E)** and Ty3 **(F)** RNA levels in WT cells and non-essential mutants of the Nup84 complex, as measured by RT-qPCR (mean±SD, n≥3, relative to WT and normalized to *ACT1* mRNAs). **(G)** Analysis of log2 fold change (FC) of expression of Ty1, Ty2 and Ty3 copies in *nup133Δ (top panel)* or *nup84Δ (bottom panel)* mutants relative to WT cells. Only reads assigned to a single locus in the genome were considered for this analysis. **(H)** Scheme of the Ty1-*lacZ* reporter. The *lacZ* gene is fused in-frame to *GAG* at individual endogenous Ty1 copies downstream of their cognate promoter sequences (at coordinate 1571 of Ty1-H3) [55]. The position of the qPCR amplicon used in (I) and (J) is indicated. Scale bar, 50bp. **(I)** Ty1-*lacZ* fusion RNA levels in WT cells, as measured by RT-qPCR (mean±SD, n=3, relative to DR1 and normalized to *ACT1* mRNAs). The Ty1 copies are ranked according to their RNA levels and annotated as highly or weakly expressed in [55]. **(J)** Ty1-*lacZ* RNA levels of different Ty1-*lacZ* fusions, as measured by RT-qPCR in WT cells and *nup133*Δ or *nup84*Δ mutants (mean±SD, n=3, relative to WT and normalized to *ACT1* mRNAs). ns, not significant; * p<0.05; ** p<0.01; *** p<0.001; **** p<0.0001, two-sided Wilcoxon rank-sum test (panels **C,G**) or Welch’s t test with comparison to the WT strain (**D,E,F,J**). Note that the low number of Ty3 loci precludes any statistical analysis (**C,G**).

To determine whether this novel function of Nup133 and Nup84 is shared by the other components of the Nup84 complex, we measured Ty1, Ty2 and Ty3 RNA levels by RT-qPCR in deletion mutants for all non-essential Nups of the Nup84 complex in the S288C genetic background. As a control for the ability of our RT-qPCR measurement to distinguish Ty families, deletion of *SPT3,* encoding a SAGA subunit specifically required for Ty1 transcription, strongly reduced Ty1 RNA levels while it did not affect Ty3 expression (Fig S1C). Our assay further scored a significant ~2-fold increase in Ty1 RNA levels in four of the five tested Nup84 complex mutants (Fig 1D). Similarly, Ty2 and Ty3 RNA levels increased in all the mutants except *seh1*Δ (Fig 1E-F), in line with the less severe phenotypes displayed by this deletant as compared to other Nup84 complex mutants [54]. Altogether, these data highlight an unexpected function of the Nup84 complex in limiting the accumulation of RNA from LTR-retrotransposons.

Dozens of Ty copies are scattered throughout the yeast genome and could be differently affected by NPCs. To address whether the Nup84 complex specifically targets certain Ty within the genome or globally regulates the expression of all the LTR-retrotransposon copies, we performed a differential expression analysis based on single-mapping reads to discriminate the expression changes of the distinct genomic Ty loci (Fig 1G). About 80% of the reads multi-mapping to Ty1 and Ty2 copies and 30% corresponding to the two Ty3 copies were thus filtered out. However, the differential expression analysis based on the remaining Ty reads gave fully consistent results with the multi-mapping read analysis, with similar median log2 fold changes for Ty1 copies (single-mapping: *nup133Δ:* 0.43, *nup84Δ:* 1.02; multimapping: *nup133Δ:* 0.46 and *nup84Δ:* 1.2; compare Fig 1G and 1C). Moreover, the single-mapping analysis showed that all the 32 genomic copies of Ty1 were up-regulated in both *nup133*Δ and *nup84*Δ mutants, although to different extent depending on the copies (Fig 1G). Similarly, the expression of the two Ty3 copies was highly up-regulated in both mutants of the Nup84 complex. In contrast, Ty2 copies were all up-regulated in the *nup84*Δ mutant while some of them were not affected in the *nup133*Δ mutant, explaining the lower overall impact of the *nup133*Δ mutant compared to the *nup84*Δ mutant on Ty2 transcript levels (Fig 1C).

The estimation of the transcript levels derived from different TE genomic copies remains partially biased with such transcriptomic analyses. In particular, single-mapping will retain more reads on older sequences that have accumulated more SNPs than younger elements [41]. To precisely evaluate the global impact of the Nup84 complex on distinct copies of Ty1, we used integrated Ty1-*lacZ* fusions previously designed to study the transcription of individual Ty1 copies at their genomic location [55]. In these constructions, the *lacZ* gene was fused at position 1571 of each Ty1 copy to keep an intact Ty1 promoter that extends over 1kb from the 5’LTR to a large part of the *GAG* ORF (Fig 1H). RT-qPCR measurement of the RNA levels produced from representative Ty1-*lacZ* copies revealed a range of expression levels in WT cells, in agreement with earlier β-galactosidase activity measurements [55] (Fig 1I). Importantly, *Ty1-lacZ* RNA levels increased for every Ty1 copy tested upon deletion of *NUP133* and *NUP84* (Fig 1J), similarly to the overall accumulation of RNAs produced by all the endogenous Ty1 copies of these strains (Fig S1D), and in agreement with our single-read mapping analysis. While slight variations in the upregulation levels were noticeable between Ty1-*lacZ* fusions, they did not correlate with the basal expression of the Ty1 copy in WT cells (Fig 1I), its position on the Watson or Crick strand or on the chromosome arm, or the close proximity of a *tDNA* gene (Fig S1E). Taken together, these results demonstrate that the Nup84 complex broadly restricts the expression of Ty LTR-retrotransposons all over the genome.

### The Nup84 complex regulates the transcription of Ty LTR-retrotransposons

An increase in RNA levels could stem from changes in transcription or RNA stability. Since Ty1 RNAs have been described as particularly stable [56] and considering that mutations of the Nup84 complex have already been associated with changes in transcription of reporter genes [34,35], we analyzed the influence of the Nup84 complex on Ty1 transcription. For this purpose, we used Ty1-*lacZ* fusions to analyze the recruitment of RNAP II on individual Ty1 copies by ChIP-qPCR (Fig 2A, *top panels*). In WT cells, RNAP II occupancy was higher at the *DR4-lacZ* locus than at the *LR1-lacZ* locus, in agreement with their status of highly- and weakly-expressed Ty1 copies, respectively (Fig 2A, *top panels* and 1I) [55]. Upon deletion of either *NUP133* or *NUP84*, RNAP II levels increased at both loci and on all genomic Ty1 elements, while they were unchanged at a highly-transcribed control gene (*PMA1*; Fig 2A, *top and middle panels*). Furthermore, while Ty2 and Ty3 copies were undetectably recruiting RNAP II in WT cells, significant RNAP II ChIP signals were scored at these loci upon deletion of the subunits of the Nup84 complex (Fig 2A, *lower panels*). This set of results demonstrates that the Nup84 complex globally represses the transcription of Ty1, Ty2 and Ty3 elements. While Ty1 and Ty2 are closely related LTR-retrotransposons, Ty3 belongs to another superfamily, suggesting that the Nup84 complex keeps transcription of evolutionarily-distant LTR-retrotransposons at low levels.

**Figure 2:**
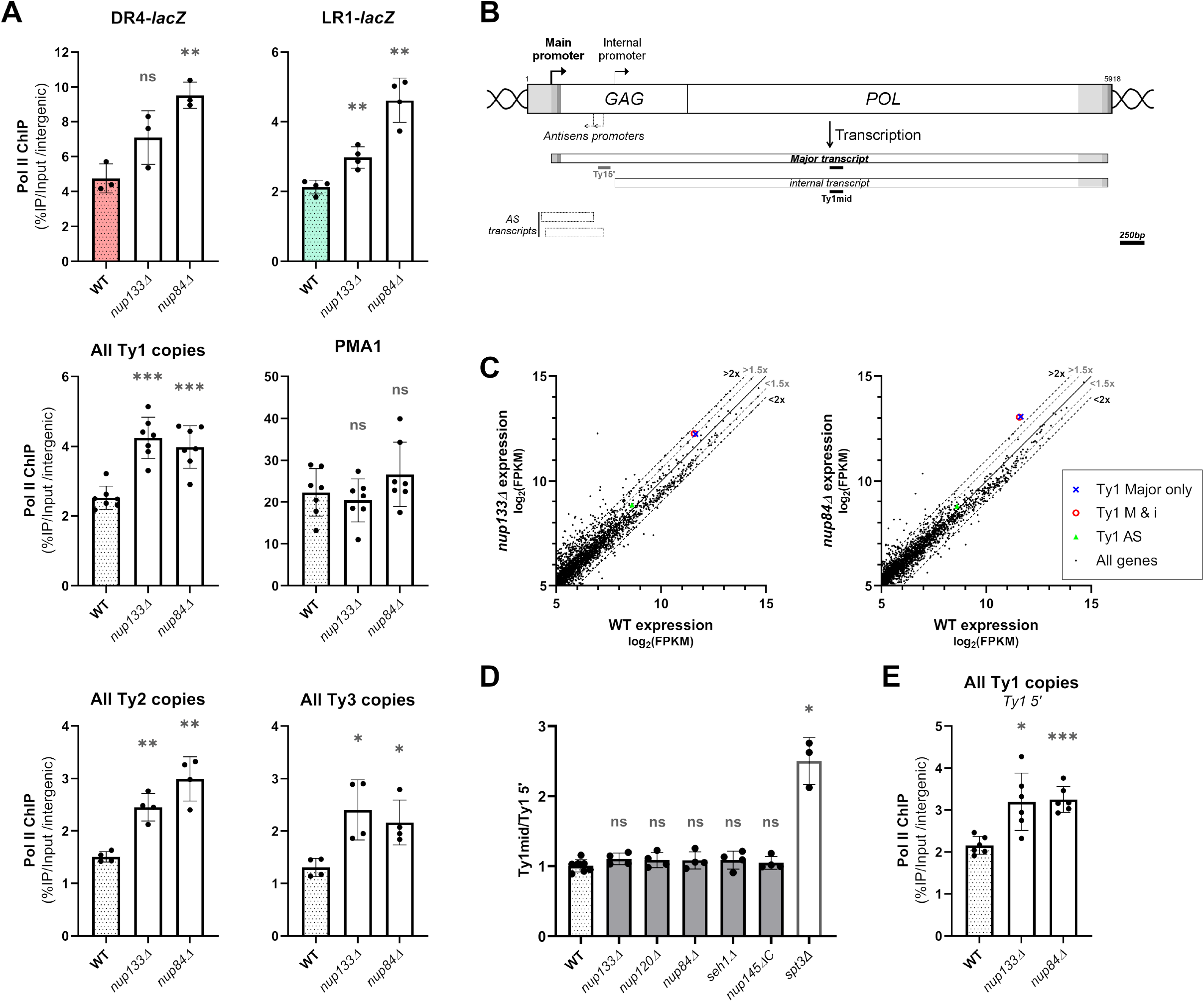
The Nup84 complex regulates the expression of LTR-retrotransposons at the transcriptional level. **(A)** RNAP II occupancy on the DR4 and LR1 Ty1-*lacZ* fusions (*top panels*), all Ty1 genomic copies and *PMA1* locus (*middle panels*), and all Ty2 and Ty3 genomic copies (*lower panels),* as determined by ChIP in WT, *nup133*Δ and *nup84*Δ cells. Values (mean±SD, n≥3) are expressed as a percentage of IP and normalized to an intergenic region. **(B)** Ty1 locus organization and expression. Long Terminal Repeats (LTR) flank two overlapping ORFs (*GAG* and *POL).* The 5.7 kb Ty1 major transcript starts at the U3/R junction of the 5’ LTR and ends at the R/U5 junction of the 3’ LTR (U3, R and U5 regions are indicated from light grey to dark grey). The 5 kb internal Ty1 RNA species is transcribed from an internal promoter located around 800bp downstream of the main promoter. Two major described species of antisense (AS) noncoding RNAs are indicated, as well as the position of the qPCR amplicons used in (**D**). The Ty1mid amplicon is used for Ty1 RNA levels measurement by RT-qPCR in all the figures except Fig 2D-E. Scale, 250bp. **(C)** Scatterplots showing expression levels of cellular RNAs in *nup133Δ (left)* or *nup84Δ (right)* mutants vs WT strains as described in Fig 1B. Expression of Ty1 based on reads either uniquely assigned to major transcripts (*blue cross*) or assigned to both major and internal transcripts (*red circle*) is specified. The abundance of antisense Ty1 RNAs is also highlighted (green triangle). **(D)** Ty1 RNA levels in WT cells and non-essential mutants of the Nup84C, as measured by RT-qPCR using Ty1 5’ and Ty1mid primers, which hybridize only with major transcripts and all sense transcripts, respectively. The ratio of values obtained with Ty1mid and Ty1 5’ is represented (mean±SD, n≥3, relative to WT). The *spt3*Δ mutant is used as a positive control for induction of the internal promoter and specific increase of internal transcripts. **(E)** RNAP II occupancy at the 5’ end of Ty1 genomic copies determined by ChIP using the Ty1 5’ amplicon in WT, *nup133*Δ and *nup84*Δ cells. Values (mean±SD, n≥3) are expressed as a percentage of IP and normalized to an intergenic region. ns, not significant; * p<0.05; ** p<0.01; *** p<0.001, Welch’s t test with comparison to the WT cells.

Different promoters have been identified in Ty1, yielding sense and antisense Ty1 RNA species (Fig 2B) (reviewed in [45]). While only the major transcript contains all the sequences required for reverse transcription, internal and antisense transcripts have been assigned roles in the regulation of retrotransposition. In particular, the internal transcript encodes p22, a truncated Gag protein that is necessary for copy number control of Ty1 mobility [57]. Although internal transcripts are weakly expressed in WT cells, this internal promoter is specifically induced in certain situations (e.g. *spt3* mutants [57]). To determine precisely which of the promoter(s) located within Ty1-*lacZ* fusions are repressed by the Nup84 complex, we first re-analyzed our RNA-seq data by separately quantifying reads assigned to three different Ty1 regions, as previously described [58]: the 5’ part of the coding strand specific to the major transcript (“Ty1 Major only”; Fig 2C), the 3’ part of the coding strand shared by both the major and the internal transcripts (“Ty1 M & i”, Fig 2C), and the 3’ part of the non-coding strand corresponding to antisense RNAs (“Ty1 AS”, Fig 2C). Read counting indicated that antisense RNA levels remained unchanged in the mutants compared to WT cells, whereas both “Ty1 Major only” and “Ty1 M & i” transcripts were equally abundant in WT cells and similarly over-represented in *nup133*Δ and *nup84*Δ mutants (Fig 2C). This suggests that the internal transcripts are produced at very low levels in all strains and that only the transcription of the major transcript is increased in the mutants of the Nup84 complex. RT-qPCR analysis comparing the amount of RNAs measured by two different qPCR amplicons in the “Ty1 Major only” or “Ty1 M & i” regions confirmed that unlike in the control *spt3*Δ mutant, the internal promoter is not induced in any of the Nup84 complex mutants (Fig 2D). The increased recruitment of RNAP II at the 5’ end of Ty1 upon deletion of *NUP133* and *NUP84* (Fig 2E) further confirmed the induction of the main promoter and strongly suggests that the Nup84 complex restricts the transcription of Ty1 at the initiation level.

Altogether, these results demonstrate that the Nup84 complex represses the transcriptional activity of Ty1, Ty2 and Ty3 LTR-retrotransposons, thereby limiting the quantity of transcripts that serve as templates for both translation and reverse transcription.

### The Nup84 complex represses Ty1 expression through Ulp1-dependent SUMOylation processes

We next asked by which mechanisms the Nup84 complex could repress Ty transcription. Although this regulation occurs for different families of retrotransposons, we focused our study on Ty1 transcripts, which are far more abundant than Ty2 and Ty3 RNAs in WT cells (about 10-fold and 600-fold, respectively; Fig 1B). We first investigated whether the derepression of Ty1 transcription was specific of mutants of the Nup84 complex, or could be generally observed when NPC structure or nucleocytoplasmic transport are compromised. For this purpose, we systematically assayed Ty1 expression by RT-qPCR in representative nucleoporin mutants with characterized defects in the NPC permeability barrier (*nup170Δ, nup188*Δ, [59]), NLS-dependent protein import (*nup2*Δ, [60]) or tRNA export (*nup100*Δ, [61]) (Fig 3A). None of these mutants displayed altered Ty1 RNA levels as compared to WT, in contrast with the *nup133*Δ mutant (Fig 3B), supporting the fact that Ty1 derepression is not the mere consequence of impaired nucleo-cytoplasmic transport in Nup84 complex mutants.

**Figure 3:**
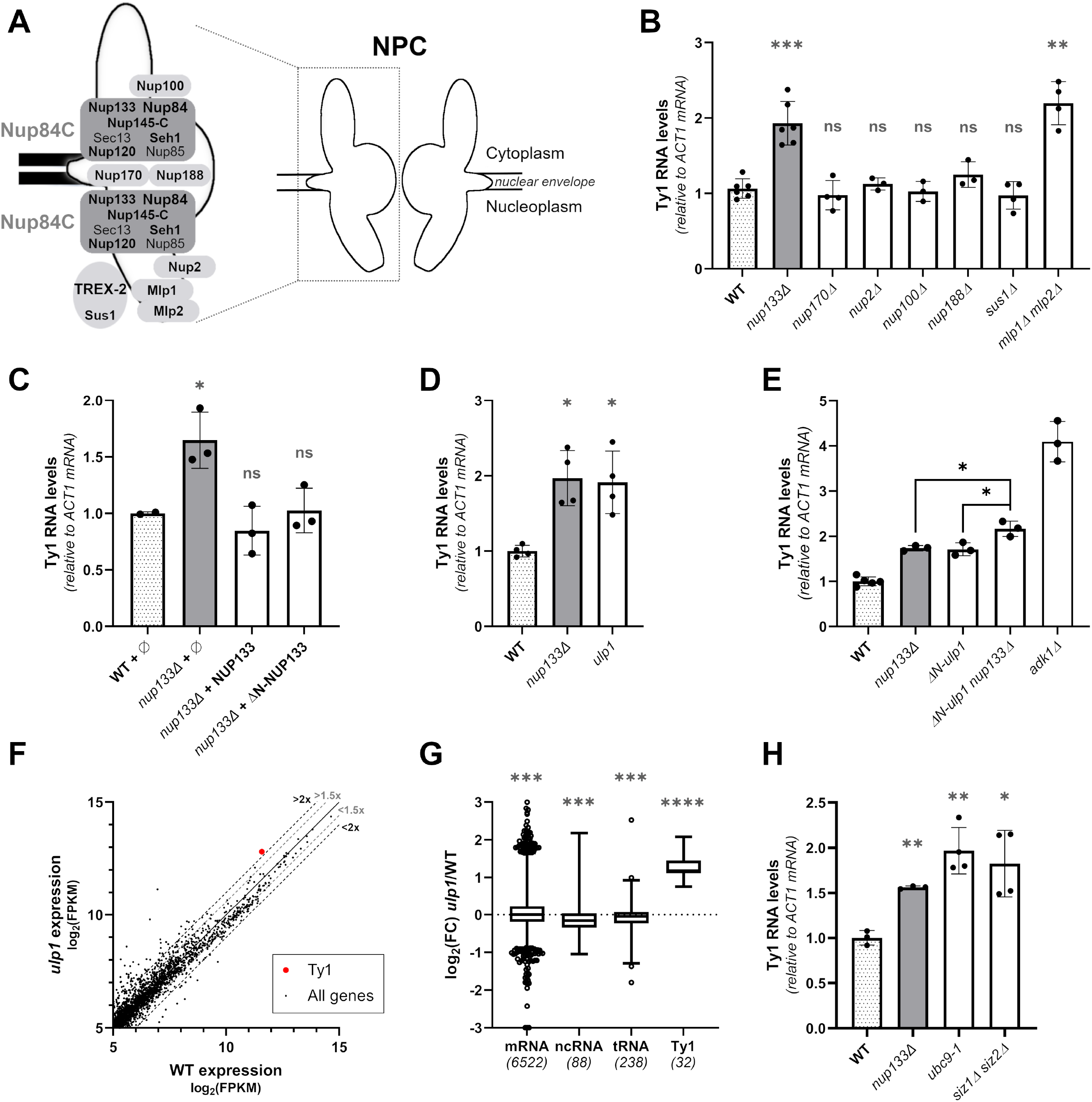
The Nup84 complex represses Ty1 expression through the tethering of the SUMO protease Ulp1 at the NPC. **(A)** Schematic representation of the structural organization of the NPC, highlighting the relative position of the nucleoporins analyzed in this study. Nup84 complex (Nup84C) subunits and other nucleoporins mutated here appear in bold. **(B)** Ty1 RNA levels in WT cells and mutants of nucleoporins and NPC-associated proteins, as measured by RT-qPCR (mean±SD, n≥3, relative to WT and normalized to *ACT1* mRNAs). Ty1 RNA levels in the *nup133*Δ mutant are shown as a control of Ty1 induction. **(C)** Ty1 RNA levels in *nup133*Δ cells carrying either the pUN100 (ø), the pUN100-protA-NUP133 or the pUN100-protA-ΔN-NUP133 plasmids, as measured by RT-qPCR (mean±SD, n≥2, relative to WT and normalized to *ACT1* mRNA values). **(D)** Ty1 RNA levels in WT cells and the *ulp1* mutant, as measured by RT-qPCR (mean±SD, n≥3, relative to WT and normalized to *ACT1* mRNAs). Ty1 RNA levels in the *nup133*Δ mutant are shown as a control of Ty1 induction. **(E)** Ty1 RNA levels in WT cells and single or double *nup133Δ ΔN-ulp1* mutants, as measured by RT-qPCR (mean±SD, n≥3, relative to WT and normalized to *ACT1* mRNA). Ty1 RNA levels in the *adk1*Δ mutant are shown as a control of strong Ty1 induction. **(F)** Scatterplot showing expression levels of cellular RNAs in *ulp1* mutants vs WT cells as described in Fig 1B. The abundance of total Ty1 RNAs is highlighted with a red dot. The thresholds of 1.5 or 2-fold change between the 2 strains are indicated by a grey or dark dashed line, respectively. **(G)** Boxplot analysis of log2 fold change (FC) of different categories of transcripts in *ulp1* mutant relative to WT cells. The number of transcripts expressed in both strains and considered in each category is indicated in brackets. Note that the low number of Ty3 loci precludes any statistical analysis. **(H)** Ty1 RNA levels in WT cells and *ubc9-1* or *siz1Δ siz2*Δ mutants, as measured by RT-qPCR (mean±SD, n≥3, relative to WT and normalized to *ACT1* mRNAs). ns, not significant; * p<0.05; ** p<0.01; *** p<0.001, **** p<0.0001, Welch’s t test with comparison to WT (**B,C,D,H**) or as indicated **(E)**, or two-sided Wilcoxon rank-sum test (**G**).

We then examined if one of the previously described roles of the Nup84 complex in mRNA export and NPC distribution could underlie its impact on Ty1 transcription [29]. Although alterations in mRNA export can impact transcription [62], independently impairing mRNA export through the inactivation of the NPC-associated mRNA export complex TREX-2 (*sus1*Δ) did not impact Ty1 expression levels (Fig 3B). Likewise, expression of a separation-of-function *nup133* allele, which solely exhibits the NPC clustering phenotype (*ΔN-nup133;* [63]), fully complemented Ty1 RNA expression defects (Fig 3C). Altogether, our data indicates that neither the mRNA export defects nor the NPC clustering observed in Nup84 complex mutants are the cause of Ty1 derepression.

Strikingly, the increase in Ty1 RNA levels observed in Nup84 complex mutants was fully phenocopied by the simultaneous inactivation of both nuclear basket proteins Mlp1 and Mlp2 (*mlp1Δ mlp2*Δ, Fig 3B). Since the Nup84 complex and the nuclear basket have shared functions in the tethering of the SUMO-deconjugating enzyme Ulp1 to NPCs [30,64], we directly assayed whether *ULP1* inactivation could similarly impact Ty1 expression. RT-qPCR analyses revealed increased Ty1 RNA levels in a *ulp1* thermosensitive mutant impacting both Ulp1 NPC localization and catalytic activity (*ulp1-333,* referred to as *ulp1;* [65]) (Fig 3D), as well as in a *ulp1* truncation mutant lacking its pore targeting domain (*ΔN-ulp1,* [66]; Fig 3E). Further transcriptomic analysis of *ulp1* cells confirmed a specific increase of Ty1 RNAs, with other transcript categories, in particular RNAP II-transcribed mRNAs, being globally unaffected, as in Nup84 complex mutant cells (Fig 3F and G). To confirm that the Ty1 phenotype of Nup84 complex mutants originates from defective Ulp1 tethering, we performed epistasis experiments by combining both *nup133*Δ and *ΔN-ulp1* mutations. Importantly, the *nup133Δ ΔN-ulp1* double mutant did not display any synergistic effect as compared to the single mutant phenotypes, although our assay was not saturated as revealed by the higher Ty1 RNA accumulation detected in the *adk1*Δ mutant ([67]; Fig 3E). Altogether, our data establish that the Nup84 complex regulates Ty1 transcription through the tethering of the SUMO-deconjugating enzyme Ulp1.

Importantly, Ulp1 acts at distinct stages in the SUMOylation process, both activating SUMO precursors, an absolute prerequisite for SUMO conjugation, and removing SUMO moieties from certain targets. In addition, anchoring of Ulp1 at the NPC was proposed to prevent unscheduled deSUMOylation in the nucleoplasm [30,64,66]. To define whether the Ty1 derepression observed in *ulp1* mutants is caused by decreased SUMOylation or impaired deSUMOylation, we assayed Ty1 expression in other mutants of the SUMOylation machinery (Fig 3H). Inactivation of the unique SUMO-conjugating enzyme, Ubc9, or the simultaneous deletion of the two major SUMO ligases, Siz1 and Siz2, triggered a significant increase in Ty1 RNA levels, comparable to the one observed in Nup84 complex or *ulp1* mutants, supporting the fact that Ty1 activation originates from decreased SUMOylation of cellular factors. Taken together, our results support a model in which the Nup84 complex limits Ty1 transcription by regulating SUMOylation processes through the tethering of the SUMO-deconjugating enzyme Ulp1 at the NPC.

### The Nup84 complex-dependent control of Ty1 transcription prevents unrestrained retrotransposition

RNAs are pivotal in the retrotransposition process as they serve as templates for both the translation of structural and enzymatic proteins and for the reverse transcription into cDNAs (Fig 4A, *left panel*). To explore the consequences of the increase of Ty1 RNA levels observed in Nup84 complex mutants on Ty1 retrotransposition, we quantified the efficiency of each stage of the retrotransposition cycle in WT, Nup84 complex and *ulp1* mutant cells.

**Figure 4:**
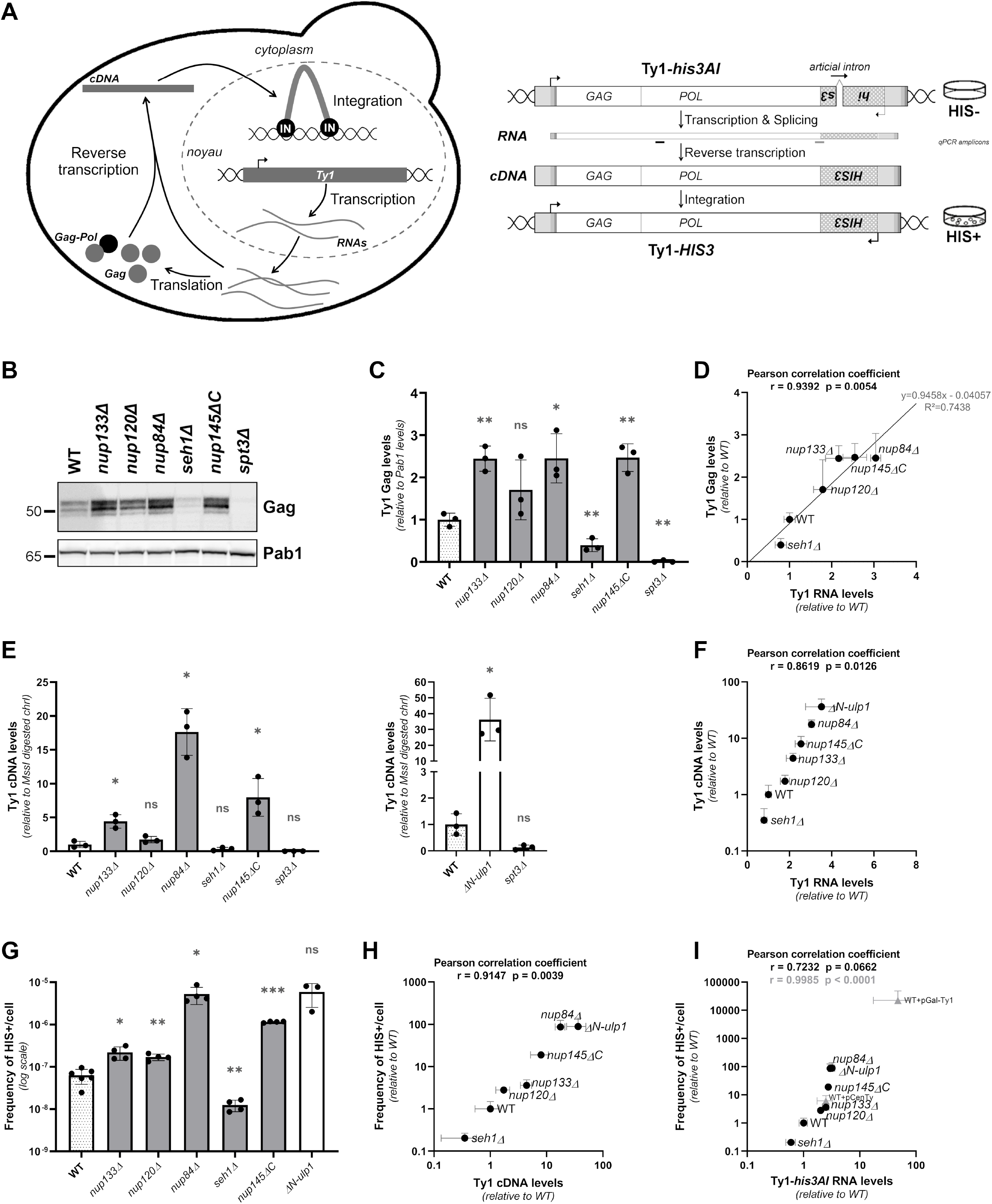
The Nup84 complex prevents unrestrained retrotransposition. **(A)** *Left panel:* The Ty1 replication cycle. A Ty1 element is first transcribed into RNAs that are exported to the cytoplasm. Ty1 RNAs are subsequently translated into Gag and Gag-Pol proteins and reverse transcribed into cDNAs by the reverse transcriptase, which is derived from the Gag-Pol polyprotein maturation into Gag, protease, integrase (IN) and reverse transcriptase proteins. IN binds the Ty1 cDNA, which is imported into the nucleus and integrated into the yeast genome. *Right panel:* Principle of the retrotransposition assay. The *his3AI* retrotransposition reporter gene consists of a *HIS3* gene inserted in the opposite orientation in the 3’ untranslated region of Ty1 and interrupted by an artificial intron, in a spliceable orientation in the Ty1 transcript. Upon splicing and reverse transcription, a Ty1-*HIS3* cDNA is produced which confers a His+ phenotype to the cells when integrated into the host genome. qPCR amplicons used for total Ty1 or specific *Ty1-his3AI* transcript quantification are indicated in black and grey, respectively. **(B)** Whole cell extracts of the indicated strains analyzed by western blotting using anti-VLP antibodies, revealing Gag proteins. Note that these antibodies detect both p49 and p45 Gag species. Pab1 is used as a loading control. Molecular weights are indicated (kDa). **(C)** Quantification of Gag levels from (B) (mean±SD, n=3, relative to WT and normalized to Pab1 protein levels). **(D)** Total Gag protein levels (mean±SD, n=3, relative to WT, values from Fig 4C) are plotted as a function of total Ty1 RNA levels (mean±SD, n≥3, relative to WT, values from Fig S2B). The Pearson correlation coefficient and associated p-value are indicated. The equation and R^2^ coefficient obtained from the linear regression are also indicated. **(E)** Total Ty1 cDNA levels in WT cells and non-essential mutants of the Nup84 complex (*left panel*) or Ulp1 (*right panel*), as measured by qPCR (mean±SD, n=3, relative to WT and normalized to the values of a chromosome I genomic locus digested by MssI). The *spt3*Δ mutant is used as a control for qPCR specificity because Ty1 expression is strongly decreased in this mutant. **(F)** Total Ty1 cDNA levels (mean±SD, n=3, relative to WT, values from Fig 4E) are plotted as a function of total Ty1 RNA levels (mean±SD, n≥3, relative to WT, values from Fig S2B) in the indicated strains. The Pearson correlation coefficient and associated p-value are indicated. **(G)** Retrotransposition frequencies (log scale, mean±SD, n≥3) of Ty1-*his3AI* reporter in WT cells and non-essential mutants of the Nup84C or *ΔN-ulp1* mutant. **(H)** Retrotransposition frequencies (mean±SD, n≥3, relative to WT, data from Fig 4G) are plotted as a function of total Ty1 cDNA levels (mean±SD, n=3, relative to WT, values from Fig 4E) in the indicated strains. The Pearson correlation coefficient and associated p-value are indicated. **(I)** Retrotransposition frequencies (mean±SD, n≥3, relative to WT, data from Fig 4G) are plotted as a function of Ty1-*his3AI* RNA levels (mean±SD, n≥3, relative to WT, values from Fig S2D) in the indicated strains. The Pearson correlation coefficient and associated p-value are indicated in black. Retrotransposition frequencies and Ty1-*his3AI* RNA levels in WT cells expressing a Ty1-*his3AI* reporter from pCenTy (centromeric) or pGAL-Ty1 (multicopy) plasmids are represented in grey. The Pearson correlation coefficient and associated p-value calculated upon inclusion of the plasmid values are indicated in grey. Note that all the strains used in this figure carry a chromosomal Ty1-*his3AI* reporter. ns, not significant; * p<0.05; ** p<0.01, *** p<0.001, Welch’s t test with comparison to the WT strain.

First, to assess the export and the translation of Ty1 RNAs, we evaluated the levels of the Ty1-encoded structural Gag protein by western blot analysis. Increased Gag protein levels were detected in all Nup84 complex mutants except *seh1*Δ (Fig 4B-C), as well as in *ulp1* mutant cells (Fig S2A), mirroring Ty1 RNA accumulation (Fig S2B), and further confirming that Ty1 RNAs are normally exported to the cytoplasm. The linear correlation between RNA and Gag variations (Fig 4D; Pearson correlation coefficient r=0.9392; α~1) further indicates that the increase in Ty1 transcription in Nup84 complex mutants equivalently impacts Gag protein levels.

Next, we examined the reverse-transcription of Ty1 mRNAs in the same mutants. To accurately monitor Ty1 cDNA variations, we adapted a strategy previously developed to quantify cDNA from retrotransposons in plants [68] (see Material and Methods). In this assay, adapters are ligated to non-integrated Ty1 cDNA molecules (that unlike genomic Ty1 copies, display free ends), allowing specific qPCR-based quantification. While as anticipated, no cDNA molecules were detected in the *spt3*Δ mutant [69], validating that our assay does not detect Ty1 genomic copies (Fig 4E), we quantified about 0.1 molecule of Ty1 cDNA per WT haploid genome (Fig S2C), a value in agreement with previous reports [70,71]. Given the abundance of Ty1 RNAs in the cellular transcriptome (Fig 1B), this result confirms the low efficiency of the post-translational steps of Ty1 replication [72]. In Nup84 complex and *ulp1* mutants, Ty1 RNA accumulation resulted in a sharp increase in Ty1 cDNA molecules, suggesting that Ty1 transcript levels may be limiting for reverse transcription (Fig 4E). The accumulation of Ty1 RNAs therefore leads to an exponential increase in cDNA molecules through reverse-transcription in Nup84 complex and *ulp1* mutants (Fig 4F), as exemplified by the *nup84*Δ mutant, in which a 3-fold increase in RNA levels results in an 18-fold increase in cDNA molecules.

To finally determine the consequences of increased Ty1 RNA levels on Ty1 retrotransposition frequency in NPC mutants, we used a chromosomal Ty1-*his3AI* reporter conferring histidine prototrophy to cells having undergone a novel retrotransposon insertion event (Fig 4A, *right panel*) [73]. Ty1-*his3AI* RNA levels were increased in all *nup* and *ulp1* mutants as compared to WT cells (except *seh1Δ;* Fig S2D), confirming that the Ty1-*his3AI* reporter is targeted by the Nup84 complex as all the endogenous Ty1 copies (Fig S2B). Consistent with previous quantifications of Ty1 RNA, protein and cDNA levels, the retrotransposition frequency significantly increased in all Nup84 complex and *ulp1* mutants, except *seh1*Δ (Fig 4G). As expected, cDNA levels and retrotransposition frequencies were strongly correlated in the different mutants (Pearson correlation coefficient r=0.9147) (Fig 4H). However, similarly to the amplification scored above at the reverse-transcription stage, variations in cDNA levels markedly resulted in even more important differences in retrotransposition frequencies in Nup84 complex and *ulp1* mutant cells. For example, in the *nup84*Δ mutant, the 18-fold increase in cDNA molecules triggered by a 3-fold RNA accumulation led to an 86-fold increase in the frequency of retrotransposition. Although a strong correlation between RNA levels and retrotransposition frequencies was still observed in all the strains (Pearson correlation coefficient r=0.7232) (Fig 4I), our results establish that the Ty1 RNA accumulation triggered by Nup84 complex deficiencies is amplified at both the reverse-transcription and the retrotransposition stages.

Our data could reflect the fact that the Nup84 complex and Ulp1 also act on the Ty1 cycle in a post-transcriptional manner, or that any increase in Ty1 mRNA levels would be similarly amplified at the retrotransposition stage. To distinguish between both possibilities, we specifically interfered with Ty1 RNA levels in WT cells by using *Ty1-his3AI* reporters carried either by a centromeric, low copy plasmid, under the control of its own promoter (pCenTy), or by a multicopy plasmid, under the control of the strong inducible *GAL1* promoter (pGAL-Ty1). Strikingly, the retrotransposition frequencies of these reporters were correlated with the differences in their Ty1-*his3AI* RNAs levels, and aligned on the correlation established for Nup84 complex and *ulp1* mutants (Pearson correlation coefficient r=0.9985) (Fig 4I). In contrast, mutants with characterized post-transcriptional alterations of Ty1 dynamics (*e.g. rrm3Δ, fus3*Δ, [71,74]) did not fit within our correlation between Ty1 RNAs levels and retrotransposition rates (Fig S2E). Altogether, these data demonstrate that a small derepression of Ty1 transcription can result *per se* in a dramatic deregulation of Ty1 retrotransposition, highlighting the essential role of the Nup84 complex in keeping retrotransposon under tight transcriptional control.

## DISCUSSION

Multiple systematic studies for regulators of the replication of LTR-retroelements in budding yeast had previously uncovered several NPC components [46–48], identifying in particular a post-transcriptional role for the nuclear basket [49]. In this report, we have rather focused on the contribution of a unique core subcomplex of the NPC, the Nup84 complex, to the expression and the retrotransposition of endogenous Ty elements. In the past, the expression of these repeated sequences might have been neglected in genome-wide analyses, mainly because reads that do not map to a unique locus are most often discarded. Here, our dedicated transcriptomic analysis pipeline has identified a shared function for most subunits of the Nup84 complex in the transcriptional repression of phylogenetically-distant Ty retroelements, including the most abundant and the most active Ty1, for which we have further investigated the mechanism involved in this regulation (Fig 1, 2).

Although the rest of the transcriptome seems to be largely unaffected in Nup84 complex mutants, it is likely that other physiological or stress situations could lead to Nup84 complex-dependent changes in gene expression. Indeed, mechanism-wise, the common phenotypic signatures of Nup84 complex and *ulp1* mutants, as well as our epistasis analysis, support the view that the Nup84 complex represses Ty1 transcription by controlling the activity of the SUMO-deconjugating enzyme Ulp1 (Fig 3, 4), which is itself sensitive to environmental conditions, alike Ty1 (*e.g.* chemical stresses, [75–77]). In this context, the regulation of Ty1 transcription through the Nup84 complex/Ulp1 axis echoes the previously reported impact of the nuclear basket in the repression of *GAL* genes [78]. For these loci, it was shown that the mislocalization of Ulp1 from the NPC resulted in down-regulation of the SUMOylation of the *GAL* promoter-bound transcriptional repressor Ssn6, triggering unscheduled gene derepression. With the large number of transcription factors known to regulate Ty1 promoters, many of them being SUMOylated (e.g. Gcr1, Tec1, Ste12, Gcn4, Rap1, Tye7; [79] and references therein), the Ty1 derepression scored in Nup84 complex or *ulp1* mutants probably originates from a combination of SUMO-dependent regulations for multiple targets in the transcriptional machinery. Since exposure of yeast cells to DNA-damaging agents also activates the transcription of Ty1 [80–83], changes in the SUMOylation and the activity of DNA damage response factors, as reported in Nup84 complex and *ulp1* mutants [30], could similarly stem for Ty1 derepression. Still, the phenotypes we observed with other mutants involved in SUMO conjugation suggest that it is the decreased SUMOylation of such yet-to-be-identified factors that ultimately leads to the activation of Ty1.

While Ty1 derepression could be triggered by a nucleoplasmic fraction of Ulp1, the proximity of Ty1 sequences with Ulp1 SUMO-deconjugating activity bound to the NPC could activate Ty1 transcription in a spatially-regulated manner, as proposed for *GAL* genes. Furthermore, beside their impact on the activity of transcription factors, SUMOylation events could also trigger the recruitment of Ty1 elements to the NPC, as established for the inducible *GAL1-10* and *INO1* loci [78,84]. In this respect, it is striking that *tDNA* loci, which are adjacent to most Ty1 sequences in the genome due to Ty1 preferential integration close to Pol III-transcribed genes (reviewed in [85]), undergo cell-cycle dependent relocalization to the NPCs [40], and that a number of transcription factors, including the *bona fide* Ty1 activators Gcn4 and Ste12, are sufficient to promote the NPC association of their target sequences [86]. At this position, Ty1 elements may integrate the signals from various NPC-associated factors, including the Mediator co-activator complex [14], which also acts at Ty1 promoters [87], or specific partners of the Nup84 complex, as suggested by the co-evolution scored between Ty1 elements and *NUP84* sequences in *Saccharomyces* yeasts [88]. In the future, genome-wide and imaging analysis investigating the position of Ty1 elements in the nucleus will be instrumental to decipher the relationships between the expression of transposable elements and their localization with respect to NPCs.

With their specific effect on Ty1 transcription, Nup84 complex mutants provide a unique tool to decipher the relationships between Ty1 RNA production and retrotransposition (Fig 4). While Ty1 RNAs are confirmed to be very abundant RNAP II transcripts in our transcriptomic data (Fig 1), in agreement with earlier estimates [43,44], they appear to be limiting for reverse-transcription, as revealed by the exponential increase in cDNA amounts driven by RNA accumulation (Fig 4F). Solving this apparent paradox will definitely require to identify which host-specific factors limit cDNA levels in cells, from their synthesis in the cytoplasmic virus-like particles to their integration into the nuclear genome. Similarly, modest changes in cDNA amounts have substantial consequences on Ty1 mobility (Fig 4H), suggesting that cDNA molecules are also limiting for retrotransposition. This observation implies that while many controls are effective in limiting the retrotransposition of Ty1 under normal growth conditions, they can be easily overridden by activating Ty transcription, especially under adverse conditions, when retrotransposition activity contributes to the response to stress. Altogether, our results thus highlight the importance of keeping the transcription of Ty elements at a level that would prevent their excessive propagation in the genome. The pivotal role of the Nup84 complex in this process thereby adds up to the ever-growing reported mechanisms that silence TE transcription in eukaryotes, including DNA methylation, chromatin repressive marks and RNA interference [89].

## MATERIAL & METHODS

### Plasmids, yeast strains and growth

All the strains used in this study are haploid and isogenic to BY4741/2 and are listed in Supplementary Table 1. Yeast cells were grown in standard yeast extract peptone dextrose (YPD) or synthetic complete (SC) media lacking appropriate amino acids. Unless indicated, yeast cells were grown at 25°C. Growth assays were performed by spotting serial dilutions of exponentially-growing cells on solid YPD medium and incubating the plates at the indicated temperatures. All plasmids and primers used in this study are reported in Supplementary Tables 2 and 3.

### Gene expression analyses

RNA quantification and RNA polymerase II chromatin immunoprecipitation were performed as described in [76]. Briefly, total RNAs were extracted from yeast cells disrupted by bead beating and purified using the Nucleospin RNA II kit (Macherey-Nagel). For RNA-seq analysis, libraries were prepared after ribosomal RNA removal and further sequenced on an Illumina NovaSeq paired-end sequencing platform with 150 bp read length by Novogene (UK) Company Limited. For RT-qPCR analysis, total RNAs were reverse-transcribed with Superscript-II reverse transcriptase (Thermo Fisher Scientific) and cDNAs were quantified by real-time PCR with a QuantStudio 5 system using the Power Track SYBR Green Master mix (Thermo Fisher Scientific) and the primers listed in Supplementary Table 3. The amounts of the RNAs of interest were normalized relative to *ACT1* mRNA values (invariant in our transcriptomic analyses), and further set to 1 for WT cells. Unless indicated, values obtained for the Ty1mid amplicon (Fig 2B) were displayed. RNA polymerase II distribution along the genes of interest was determined by ChIP as previously reported [76]. Input and immunoprecipitated DNA amounts were quantified by realtime PCR as above.

### Western blot analysis

Total protein extraction from yeast cells was performed by the NaOH-TCA lysis method [90]. Samples were separated on 4-12% SDS-PAGE gel and transferred to nitrocellulose membranes. Western-blot analysis was performed using the following antibodies: polyclonal anti-VLP antibody ([81]; 1:10,000) and monoclonal anti-Pab1 (Abcam; 1:1000). Quantification of signals was performed using the ImageJ software.

### cDNA quantification

Ty1 cDNA quantification assay was adapted from [68]. Briefly, genomic DNA (gDNA) was phenol-extracted from exponentially-growing yeast cells after cell wall disruption by digestion with zymolyase (100T; Fisher Scientific), and further isolated by ethanol precipitation. After quantification with Qubit dsDNA BR assay kit (Thermo Fisher Scientific), 1μg of gDNA were digested by the MssI restriction enzyme to generate blunt-ended fragments, similar to Ty1 cDNA ends (FastDigest enzyme; Thermo Fisher Scientific Scientific) for 2h at 37°C in a total volume of 20μL, conditions in which a complete digestion was observed. 100ng of digested gDNA were then ligated for 1 hour at 22°C with 600ng adapters (generated by annealing oligos O-AMA158 and O-AMA159; Supplementary Table 3), in the presence of 2.5 units of T4 DNA ligase (Thermo Fisher Scientific). The reaction was then stopped by incubation at 65°C for 10min. cDNA quantification by real-time PCR was performed on a 20-fold dilution of the ligation mixture with a QuantStudio 5 system using the LC480 SYBR Green Master mix (Roche) and primers hybridizing in the adapter and Ty1 LTR (Supplementary Table 3). The cDNA amounts were normalized relative to the number of genomic copies, which were evaluated by quantifying the amount of a specific MssI-digested locus on chromosome I, and further set to 1 for WT cells.

### Retrotransposition assay

Ty1 mobility was measured as previously described in [91]. Briefly, four independent clones of strains harboring a *Ty1-his3AI* chromosomal reporter were grown to saturation for 24h at 30°C in liquid YPD. Each culture was diluted thousand-fold in 2x 1mL of liquid YPD and grown for 3 days to saturation at 25°C. Aliquots of cultures were plated on YPD (2x 100μL of a 1:20,000 dilution) and SC-HIS (2x 1ml). Plates were incubated for 4 days at 30°C and colonies were counted to determine the fraction of [HIS+] prototrophs. A retrotransposition frequency was thus calculated as the median of the ratios of number of [HIS+] cells to viable cells for each of the four independent clones. Retrotransposition frequencies were then defined as the mean of at least three medians.

### Bioinformatic analyses of RNA-seq data

Analysis was performed using Galaxy (https://usegalaxy.org/). Paired-end raw reads were trimmed by the Trimmomatic tool (version 0.38.0) [92] and mapped to SacCer3 genome using HISAT2 (version 2.1.0) [93], yielding 96-98% mapped reads. Only reads properly paired were kept for further analysis (SAMtools version1.9 [94]). Then, read counts for each genomic feature was calculated by featureCounts by assigning fractions to multi-mapping reads (version 1.6.4) [95] and differential expression analysis was performed using DESeq2 (version 2.11.40.6) [96]. To restrict analysis to single-mapping reads, the featureCounts tool was used upon filtering out reads with MAPQ score below 5.

### Statistical analysis

The following statistical tests were used: a two-sided Wilcoxon rank-sum test was used to compare the median log_2_(FC) of a dataset with a hypothetical median of 0 (Fig 1C and 3G). Alternatively, the two-sided Welch’s t test was used allowing unequal variance (other figures). * p <0.05; ** p <0.01; *** p <0.001; **** p <0.001; ns, not significant. Statistical tests were performed using GraphPad Prism version 9.0.0.

## Data availability

The RNA-seq data are available in Sequence Read Archive PRJNA686644.

## AUTHORS CONTRIBUTIONS

Conceptualization, A.B., B.P., P.L.; Methodology, A.B., B.P., P.L.; Investigation, A.B., C.C.; Formal analysis, A.B.; Visualization, A.B.; Writing, A.B., B.P., P.L.; Funding Acquisition, B.P., P.L.; Supervision, B.P., P.L.

## ACKNOWLEDGEMENTS

We are very grateful to A. Asif-Laidin, V. Doye, E. Fabre and A. Nicolas for reagents; to N. Palmic for technical help; to V. Doye in whose lab this work was initiated; to A. Babour and E. Fabre for fruitful discussions; and to V. Doye and E. Fabre for their critical reading of the manuscript.

This work was supported by intramural funding from Centre National de la Recherche Scientifique (CNRS), the Université de Paris and the Institut National de la Santé et de la Recherche Médicale (INSERM), and by grants from the Agence Nationale pour la Recherche (ANR-18-CE12-0003, to B.P., ANR-17-CE11-0025 to P.L and the initiatives d’excellence (Idex ANR-11-IDEX-0005-02) and the Labex “Who am I?” (ANR11-LABX-0071) to P.L., B.P. and A.B.), Fondation ARC pour la recherche sur le Cancer (PJA 20181208112 to B.P., and PJA 20191209703 to P.L.), Ligue Nationale contre le Cancer (Comité de Paris, to B.P.) and Inserm Apprenticeship contract (to C.C.).

---

**Supplementary Figure 1:**
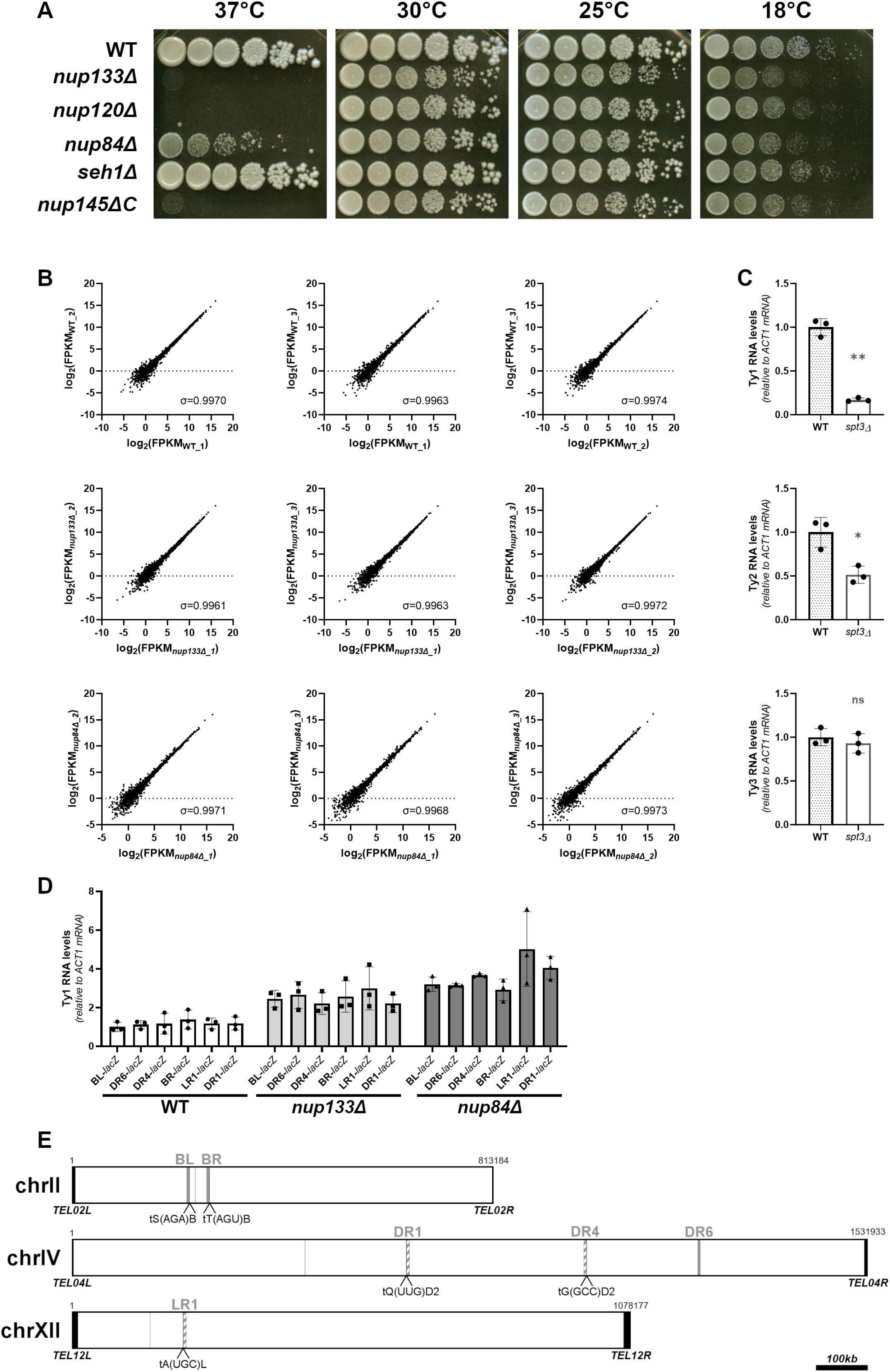
Control experiments, related to transcriptome and reporter analyses. **(A)** Characterization of the non-essential deletion mutants of the Nup84 complex. Serial 5-fold dilutions of WT and mutant cells were grown at the indicated temperatures on YPD solid medium. **(B)** Reproducibility of RNA levels measurements in RNA-sequencing data across biological replicates. RNA levels are represented as log2 (FPKM). The Spearman’s correlation coefficient is indicated for each pair of replicates. **(C)** Ty1, Ty2 and Ty3 RNA levels in WT and *spt3*Δ cells, as measured by RT-qPCR (mean±SD, n=3, relative to WT and normalized to *ACT1* mRNA values). ns, not significant; * p<0.05; ** p<0.01, Welch’s t test with comparison to the WT strain. **(D)** Total Ty1 RNA levels in the different Ty1-*lacZ* fusion strains used in Fig 1J, as measured by RT-qPCR (mean±SD, n=3, relative to WT BL-*lacZ* and normalized to *ACT1* mRNAs). **(E)** Location of the Ty1-*lacZ* fusions on chromosomes (represented to scale). Ty1 are represented by a full or hatched grey line, depending on their location on the Watson or Crick DNA strands, respectively. The presence of *tDNA*s in the close proximity of the Ty1 *-lacZ* fusions is indicated.

**Supplementary Figure 2:**
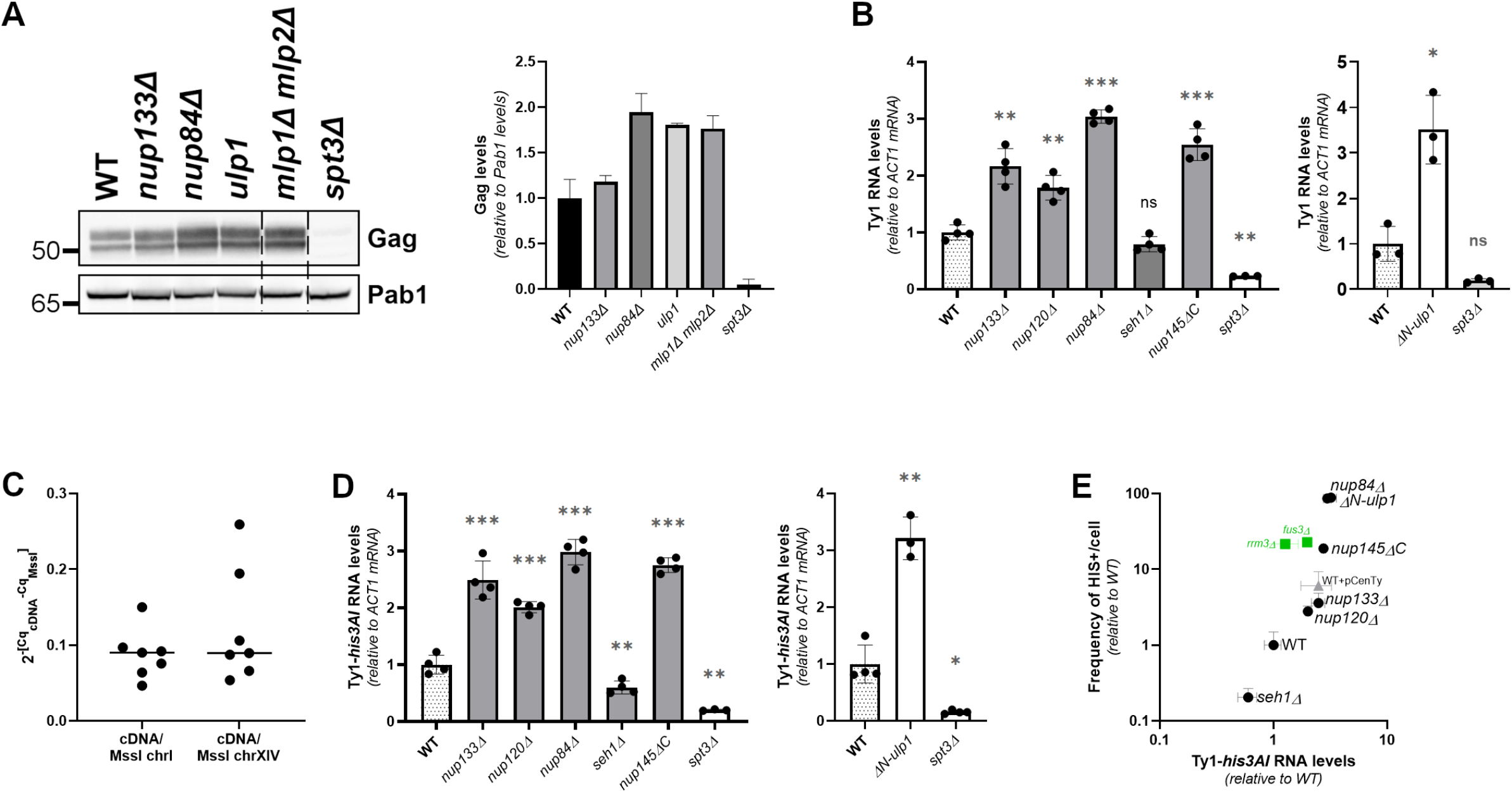
Analysis of the retrotransposition cycle in *nup* and *ulp1* mutants. **(A)** *Left panel:* Whole cell extracts of the indicated strains analyzed by western blotting using anti-VLP antibodies, revealing Gag proteins. Pab1 is used as a loading control. Molecular weights are indicated (kDa). *Right panel:* Quantification of Gag levels from western-blot analyses (mean±SD, n=2, relative to WT and normalized to Pab1 protein levels). **(B)** Total Ty1 RNA levels in WT cells and non-essential mutants of the Nup84 complex (*left panel)* or *ΔN-ulp1* mutant (*right panel),* as measured by RT-qPCR (mean±SD, n≥3, relative to WT and normalized to *ACT1* mRNAs). The *spt3*Δ mutant is used as a control for qPCR specificity because Ty1 expression is strongly decreased in this mutant. **(C)** Comparison of Ct values obtained from qPCR amplicons detecting Ty1 cDNA molecules and a genomic locus of the chromosome I or the chromosome XIV previously digested by MssI. Values and medians of 7 different experiments are plotted. **(D)** Ty1-*his3AI* RNA levels in WT cells and nonessential mutants of the Nup84C (*left panel*) or *ΔN-ulp1* mutant (*right panel*), as measured by RT-qPCR (mean±SD, n≥3, relative to WT and normalized to *ACT1* mRNAs). The *spt3*Δ mutant is used as a control for qPCR specificity, as above. **(E)** Retrotransposition frequencies and Ty1-*his3AI* RNA level measurements (mean±SD, n≥3, relative to WT) are plotted in the indicated mutants as in Fig 4I, including *rrm3*Δ and *fus3*Δ mutants (in green). ns, not significant; * p<0.05; ** p<0.01; *** p<0.001, Welch’s t test with comparison to the WT strain.

**Supplemental Table 1.**
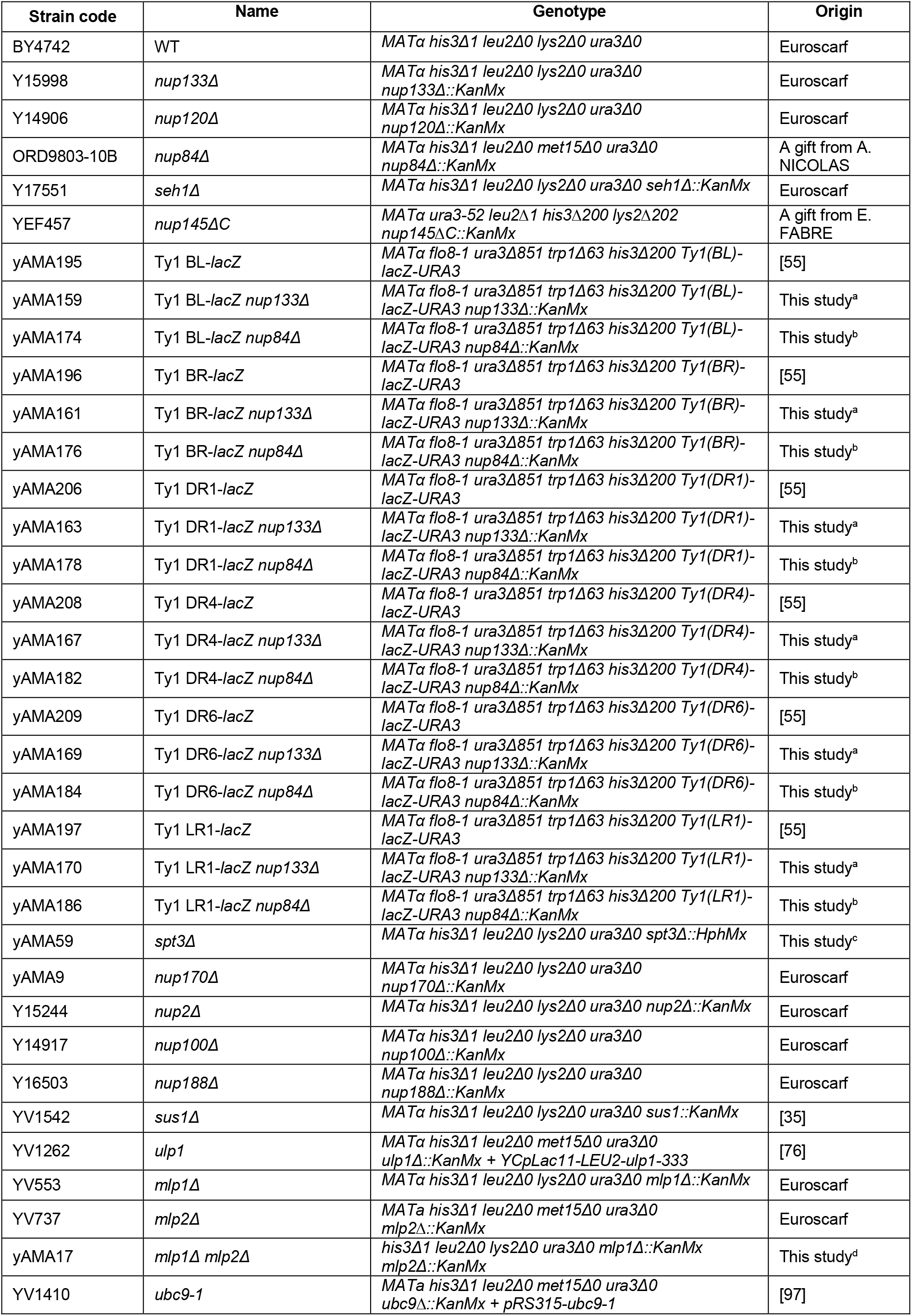
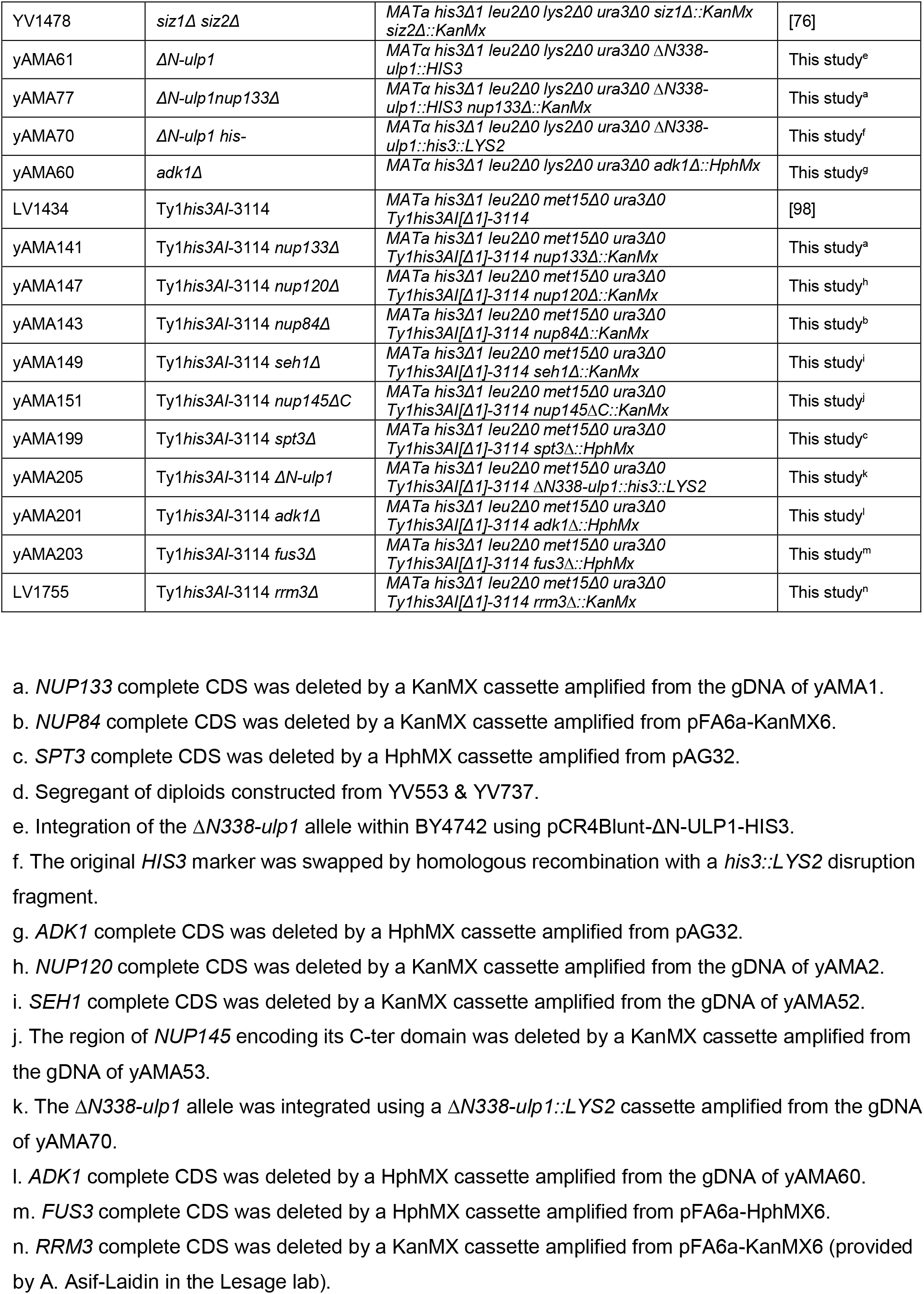
Yeast Strains used in this study.

**Supplemental Table 2.**
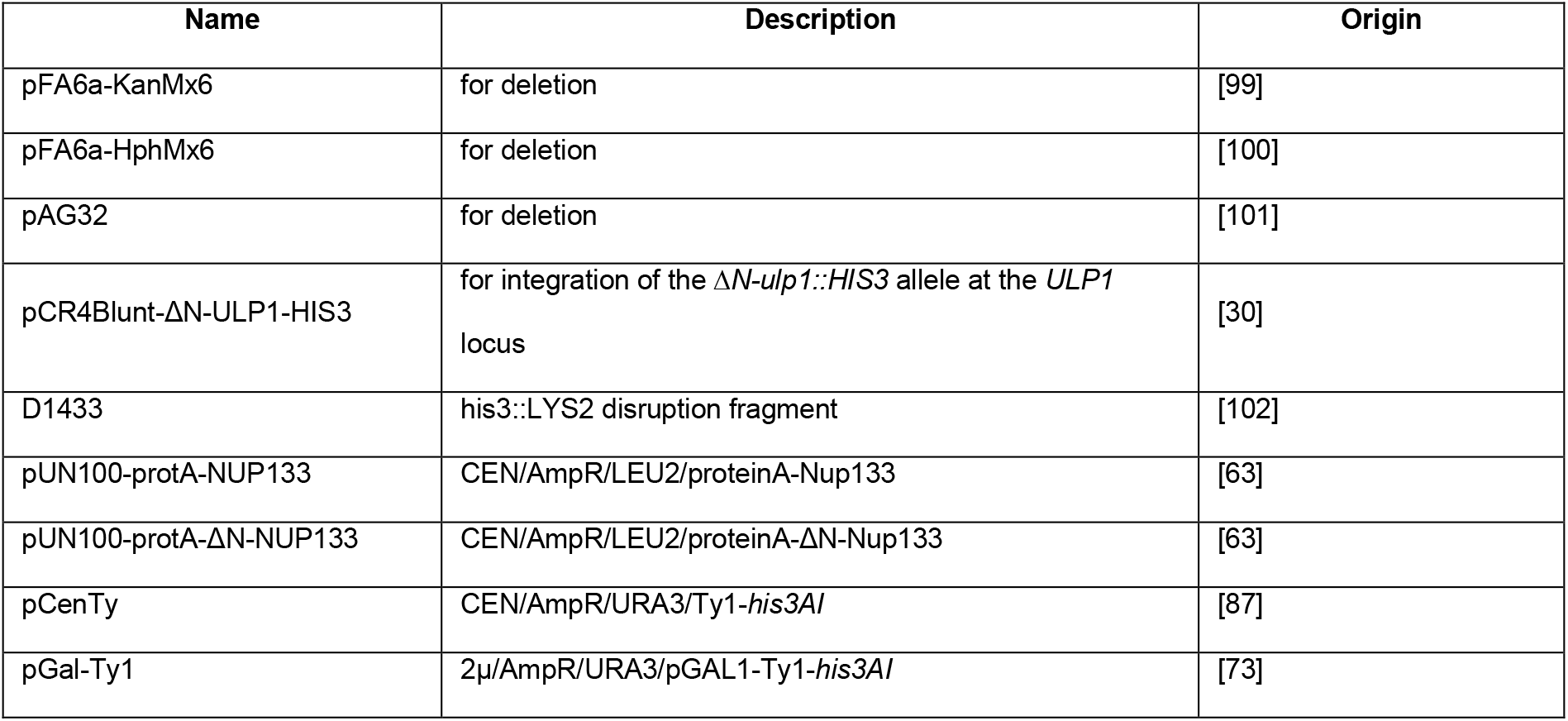
Plasmids used in this study.

**Supplemental Table 3.**
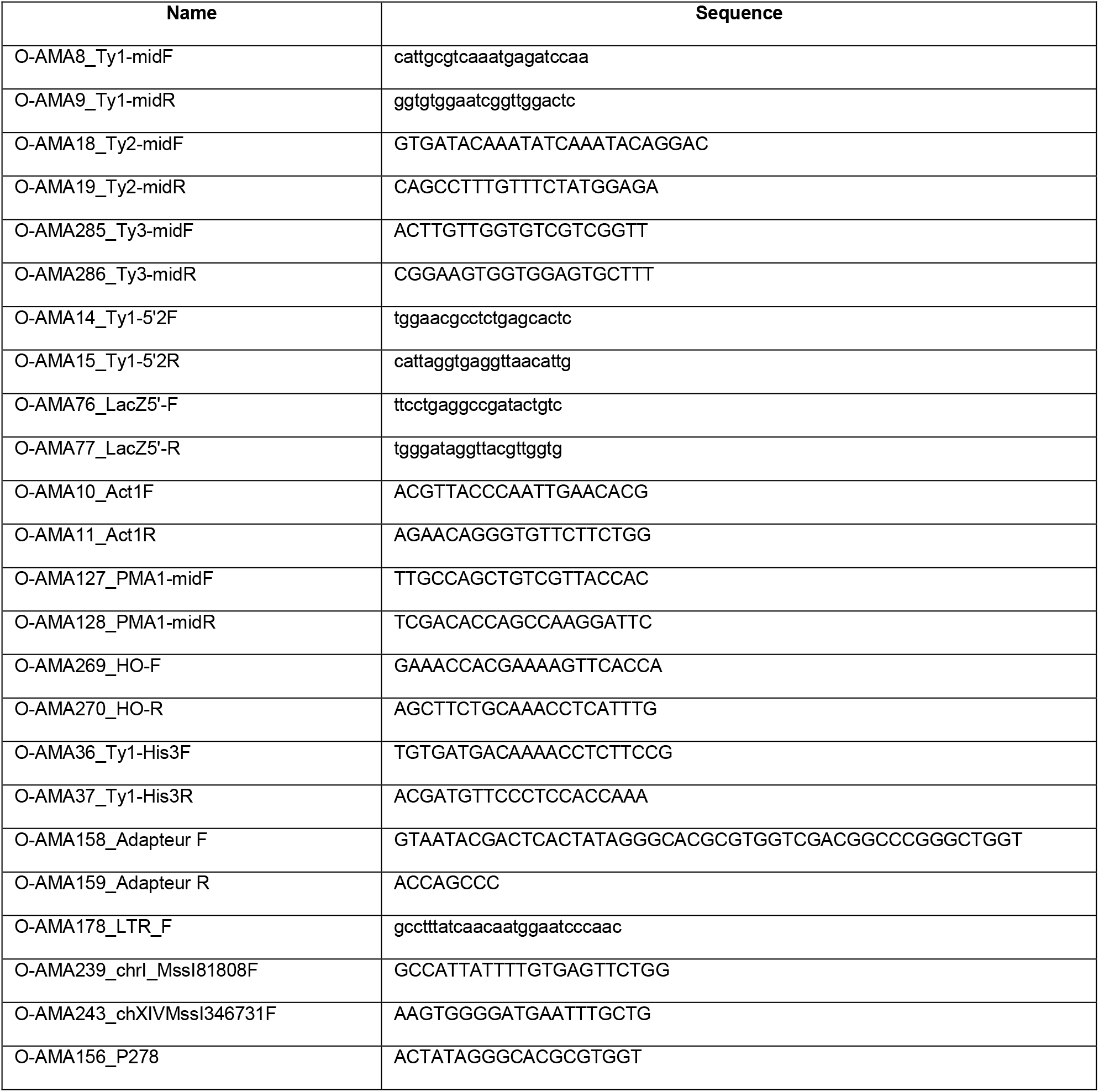
Primers used in this study.

## Notes

### Competing Interest Statement

The authors have declared no competing interest.

